# Case-control association mapping without cases

**DOI:** 10.1101/045831

**Authors:** Jimmy Z Liu, Yaniv Erlich, Joseph K Pickrell

**Affiliations:** New York Genome Center, New York, NY, USA; Department of Computer Science, Columbia University, New York, NY, USA; Department of Biological Sciences, Columbia University, New York, NY, USA

## Abstract

The case-control association study is a powerful method for identifying genetic variants that influence disease risk. However, the collection of cases can be time-consuming and expensive; if a disease occurs late in life or is rapidly lethal, it may be more practical to identify family members of cases. Here, we show that replacing cases with their first-degree relatives enables genome-wide association studies by proxy (GWAX). In randomly-ascertained cohorts, this approach enables previously infeasible studies of diseases that are absent (or nearly absent) in the cohort. As an illustration, we performed GWAX of 12 common diseases in 116,196 individuals from the UK Biobank. By combining these results with published GWAS summary statistics in a meta-analysis, we replicated established risk loci and identified 17 newly associated risk loci: four in Alzheimer’s disease, eight in coronary artery disease, and five in type 2 diabetes. In addition to informing disease biology, our results demonstrate the utility of association mapping using family history of disease as a phenotype to be mapped. We anticipate that this approach will prove useful in future genetic studies of complex traits in large population cohorts.

## INTRODUCTION

In a typical case-control genetic association study, a researcher genotypes a set of individuals that have a disease (the “cases”) and a set of individuals that do not have the disease (the “controls”). For each genetic variant, the difference in allele frequency between cases and controls can be used to estimate the causal effect of the genetic variant on the disease (assuming all potential confounders have been accounted for). While powerful, this study design requires an *a priori* decision about which disease is of interest, as well as substantial effort to identify matched cases and controls. An alternative approach is a cohort study, in which individuals are sampled from the general population and many phenotypes (along with genotypes) are collected on each individual. An advantage of a cohort study is that the cohort can be subdivided to create case-control studies of many different diseases.

However, cohort studies are limited by the fact that unbiased sampling may not yield sufficient numbers of cases to enable powerful case-control studies. For example, even in a perfectly representative sample of a million people one expects only 10,000 cases of a disease like schizophrenia with a population prevalence of 1%. Further, participants in a cohort study are rarely a fully representative sample of a population; a disease may also be rare in a cohort for the simple reason that the sampled population does not include the demographic group where the disease is most prevalent. For example, the UK Biobank (an ongoing and widely-available cohort study) sampled individuals in the age range of 40-69 (at the time of recruitment) [Sudlow et al., 2015]. By definition, this cohort does not include individuals with lethal childhood diseases, and at present there are only a handful of individuals with Alzheimer’s disease or other late-onset diseases. Similarly, cohort studies that focus on individuals of a single sex (like the Nurses Health Study) have little power to study diseases that are more common in the other sex. Other sampling approaches, like cohorts made from customers of consumer genomics companies (e.g. Eriksson et al. [2010], DNA.Land), have analogous limitations. More generally, the number of cases of a given disease present in a cohort will be a function of aspects of the disease (with rarely-occurring or rapidly-lethal diseases being more rare) and aspects of the sampling.

In this paper, we consider a study design where the researcher genotypes family members of cases, rather than cases themselves (since the cases may be difficult or impossible to contact). This design is popular in studies of longevity (where “cases” are long-lived individuals, see e.g. Barzilai et al. [2003]; Joshi et al. [2016]; Pilling et al. [2016]; Tan et al. [2010]), but has not been widely used in other situations. The approach can be thought of as taking pedigree-based association methods that allow for missing genotype data (e.g. Gudbjartsson et al. [2008]; Kong et al. [2009]; Thornton and McPeek [2007]) to an extreme where no cases have been genotyped or phenotyped by the researcher.

As a motivating example for this type of design, consider Alzheimer’s disease. As of March 25, 2016, there are 55 cases of Alzheimer’s disease listed among the approximately 500,000 individuals in the UK Biobank. However, over 60,000 individuals note that one or both of their parents was/is affected with the disease. An individual with a single affected parent can be thought to have one chromosome sampled from a population of “cases” and one from a population of “controls”. If the allele frequency (in the standard case-control setting) of some variant that increases risk of a disease is *f_A_* in cases and *f_U_* in controls, then the allele frequency in individuals with a single affected parent is
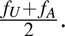.
This motivates a “proxy-case”- control association study where “proxy-cases” are the relatives of affected individuals and “controls” are the relatives of unaffected individuals. We refer to this approach as a Genome-Wide Association study by proXy (GWAX).

## Results

### Power of genome-wide association by proxy

We first explored the power of this approach with simulations and analytical calculations. Specifically, we focused on the situation where we have information about the diseases of the parents of an individual (Methods). We initially considered the case where we have no phenotype information about genotyped individuals themselves, though we consider this case later on.

The GWAX approach using proxy-cases who have one affected first-degree relative reduces the log odds ratios by a factor of around two when compared with a traditional case-control design (assuming an additive model for the impact of a genetic variant on a disease). This reduction in effect size reduces power to detect association. However, using proxy-cases may increase the effective sample size (in a cohort study) or be more logistically feasible than collecting standard cases, thus offsetting this loss in power. We calculated the number of proxy-cases and controls required such that the power to detect association is equivalent to using true cases and controls (Supplementary Note). Across the allele frequency and effect size spectrum, the proxy-case-control approach is more powerful when there are about four times (or more) as many proxy-cases and controls as there are true cases and controls, assuming the ratios of controls to cases and controls to proxy-cases are the same (Figure 1A). This ratio increases to ~ 4.9 if 10% of controls are in fact misclassified proxy-cases (Supplementary Note, Figure S8). For late onset diseases such as Alzheimer’s disease (1.6% in the population vs. 42% in those over the age of 84 [Hebert et al., 2003]) and Parkinson’s disease (0.3% in the population vs. 4% in those over 80 [de Lau and Breteler, 2006]), the proxy-case-control design gains substantial power if cohorts are sampled randomly from the population.

We next explored the situation where we have information about the phenotypes of the genotyped individuals as well. In this situation, we have “true cases” (genotyped individuals with a disease), “proxy-cases” (unaffected individuals with a parent with the disease), and “controls”. We considered analysis approaches that treat all three of these groups separately (in a 3 × 2 chi-squared test) or lumping together the “proxy-cases” and “true cases” and performing a standard 2 × 2 chi-squared test. When both true and proxy-cases are available in a population cohort study, accounting for this fact increases power (Figure 1B, Supplementary Note). For instance, for a disease with 5% prevalence and 50% heritability on the liability scale across all age groups, we expect to observe 5,000 cases and 8,597 proxy-cases in a randomly sampled cohort of 100,000. Here, for a SNP with allele frequency 0.1 in controls and an odds ratio of 1.2, there is 60.2% power at *α* = 5 × 10^−8^ to detect association using a standard 2 × 2 chi-squared test of true cases vs. controls, 87.2% power using a 2 × 2 test where cases and proxy-cases are lumped together, and 89.8% power using a 3 × 2 test where true cases, proxy-cases and controls are treated separately (Supplementary Note). In this situation, treating cases, proxy-cases and controls separately boosts the effective sample size by 1.34× when compared to a case-control design. Overall, the boost in effective sample size ranges from 1.36× to 1.28 × for disease prevalences from 1% to 20%. When disease prevalence is greater than around 34%, the test where cases and proxy-cases are lumped together is less powerful than a standard case-control test since there are no further gains in effective sample size. Nevertheless, across simulated effect sizes, allele frequencies, heritability and disease prevalences, the 3 × 2 test is consistently more powerful than the case-control test (see Supplementary Note for details).

**Figure 1.**
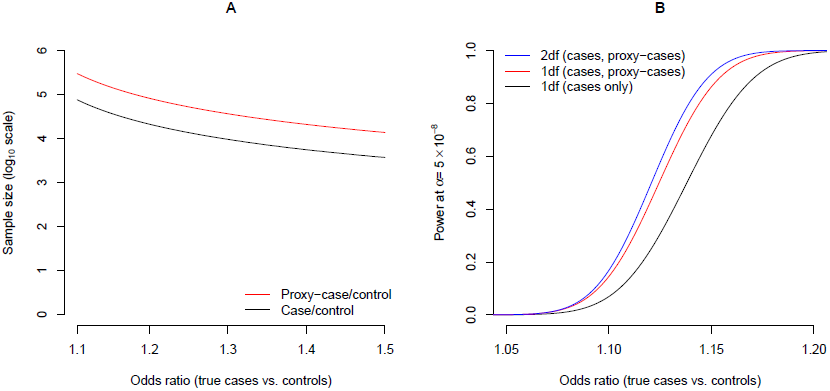
Power of proxy-case-control association designs. (A) Total sample size required for 80% power to detect association at *α* = 5 × 10^−8^ for case-control (black line) and proxy-case-control (red line) designs at a SNP with 0.1 frequency in controls. (B) Power to detect association at α = 5 × 10^−8^ using two designs that account for cases and proxy-cases (in red and blue) and a standard case-only/control design (in black). The total sample size = 100,000, disease prevalence = 0.1, heritability of liability = 0.5 and allele frequency in controls = 0.1 (See Supplementary Note).

### Application to the UK Biobank

We performed GWAX of 12 diseases in the UK Biobank (May 2015 Interim Release). After quality control and 1000 Genomes Phase 3 imputation (Methods), ~ 10.5 million low-frequency and common (*MAF* > 0.005) SNPs from 116,196 individuals of European ancestry were available for analysis. All of these individuals answered questionnaires regarding the diseases of their family members (though the medical records of the individuals themselves are available, we did not use them in this analysis in order to illustrate the approach without using cases). The number of proxy-cases per phenotype ranged from 4,627 for Parkinson’s disease to 54,714 for high blood pressure (Table S1). Based on these sample sizes, we expect greater power to detect association using GWAX than a case-control GWAS for 11 of the 12 phenotypes (high blood pressure being the exception) in the UK Biobank cohort (Figure 2).

**Figure 2.**
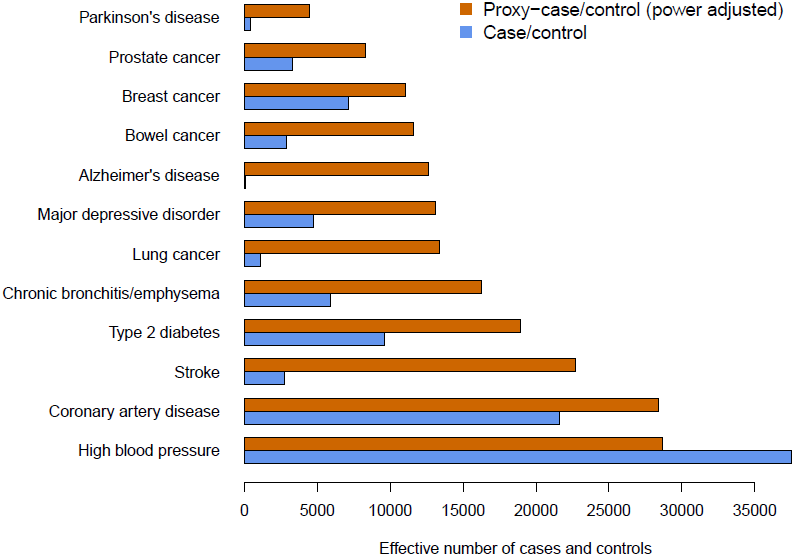
Effective sample sizes of case-control versus proxy-case-control association designs in the UK Biobank. The effective sample size for each design is: 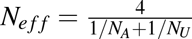 where *N*_*A*_ is the number of cases (or proxy-cases) and *N*_*U*_ is the number of controls. To account for power, we divided the effective sample size in proxy-cases/controls by four. Case/control counts for breast cancer and prostate cancer only include females and males, respectively.

Association testing was performed using logistic regression with age, sex and the first four principal components as covariates. For each SNP, we calculated an adjusted odds ratio (OR), which is directly comparable (under a standard additive model) with ORs estimated from traditional case-control study designs (Methods, Figure S1). The overall association results across the 12 phenotypes are shown in Manhattan plots, which show several clear peaks of association (Figure S2).

In the GWAX of these 12 diseases, 24 loci reached “genome-wide significance” (*P* < 5 × 10^−8^). For Alzheimer’s disease, breast cancer, heart disease, high blood pressure, lung cancer, prostate cancer and type 2 diabetes, all of these represented replications of established associations (Table S3). Among the most strongly associated loci include *APOE* (rs429358, *P* = 9.72 × 10^−195^) for Alzheimer’s disease [Corder et al., 1993], *LPA* (rs10455872, P = 2.55 × 10^−25^) and *CDKN2A/CDNKN2B* (rs4007642, P = 7.64 × 10^−21^) for coronary artery disease [Danesh et al., 2000; The Wellcome Trust Case Control Consortium et al., 2007], *FES/FURIN* (rs8027450, *P* = 6.12 × 10^−13^) for high blood pressure/hypertension [The International Consortium for Blood Pressure Genome-Wide Association Studies, 2011], *FGFR2* (rs2981583, *P* = 3.62 × 10^−12^) for breast cancer [Hunter et al., 2007], *TCF7L2* (rs34872471, *P* = 7.76 × 10^−45^) for type 2 diabetes [Grant et al., 2006], and *CHRNA5/CHRNA3* (rs5813926, *P* = 1.67 × 10^−9^ for lung cancer [Hung et al., 2008]. We identified two genome-wide significant loci for Parkinson’s disease, one of which corresponds to the established *ASH1L* locus (rs35777901, *P* = 2.25 × 10^−8^) [Nalls et al., 2014]. The second locus at *SLIT3* (rs1806840, *P* = 6.39 × 10^−9^) is implicated in Parkinson’s disease risk at genome-wide significance for the first time, although this SNP is reported as “non-significant” (P > 0.05, see URLs) in Nalls et al. [2014]. The locus remains genome-wide significant (rs1806840, *P* = 5.90 × 10^−9^) when running a linear mixed model association, suggesting that the signal is unlikely to be driven by cryptic population structure (Supplementary Note). Future genetic studies of Parkinson’s disease will be needed to determine whether *SLIT3* is a true risk locus.

### Effect size comparisons

In principle, the adjusted odds ratios obtained from a proxy-case-control design might differ from those obtained from a standard case-control design for a number of reasons. First, non-additive effects will distort these ORs in different ways in the two study designs. For example, under an additive model and Hardy-Weinberg equilibrium, the allelic odds ratio (*OR_allelic_,* estimated from allele counts) is equivalent to the heterozygote odds ratio (*OR_het_*, estimated from genotype counts), and the homozygote odds ratio (*OR_hom_*) is simply 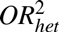. [Sasieni, 1997]. When the risk-increasing allele is partially recessive, then 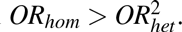. In this case, if additivity is assumed, then the observed *OR_allelic_* will be inflated by recessive effects, such that *OR_allelic_ > OR_het_* [Sasieni, 1997]. As such, the adjusted OR from GWAX (which is equivalent to *OR_het_* under additivity) will underestimate the observed *OR*_*allelic*_ from a case-control design.

Similarly, errors made by offspring in recalling the diseases of their parents would bias our estimates, as would direct causal effects of an offspring’s genotype on a parental phenotype (if, for example, a partially-heritable childhood behavior influences the diseases of their parents). Indeed, across 11 of the 12 phenotypes, females were significantly more likely to report a first-degree relative with the disease than males (Table S2), indicating at least some recall bias. Phenotype misclassification will also bias the effect size estimates. For instance, UK Biobank participants were asked whether their parents/siblings were diagnosed with “diabetes”, without any distinction between type 1 and type 2 diabetes. Given the population prevalences of type 1 and type 2 diabetes, we would expect over 90% of the proxy-cases to be type 2 diabetes. As such, we refer to this group as type 2 diabetes throughout this study.

Differences between the adjusted ORs and those previously reported may also reflect inherent differences in the samples that are collected as part of a case-control study versus a population cohort. Our additive model assumes that the frequency of a risk allele in individuals with two affected parents is the same as that in the population of cases generally. To test whether this is the case, we constructed polygenic risk scores [The International Schizophrenia Consortium, 2009] in the UK Biobank samples using previously reported ORs at established risk loci for Alzheimer’s disease [Lambert et al., 2013], coronary artery disease [The CARDIoGRAMplusC4D Consortium, 2015] and type 2 diabetes [DIAbetes Genetics Replication And Meta-analysis (DIAGRAM) Consortium et al., 2014]. Dividing the UK Biobank individuals into those affected with disease and those unaffected with two affected parents, we found significant differences in the mean polygenic risk scores between the two groups across all three disorders (*P* < 0.003, Figure S3). These results may reflect non-additive effects, or alternatively may represent a true difference in polygenic risk between the two groups. That is, given that these disorders generally occur later in life, cases ascertained as part of a case-control study (or UK Biobank participants who are aged under 69) may represent a more extreme version of the disease, harboring a greater burden of risk variants than cases that are truly sampled randomly from the general population.

To test the extent of these biases, we obtained summary association statistics from previously published GWAS for four phenotypes: Alzheimer’s disease [Lambert et al., 2013], coronary artery disease [The CARDIoGRAMplusC4D Consortium, 2015], major depressive disorder [Ripke et al., 2013] and type 2 diabetes [DIAbetes Genetics Replication And Meta-analysis (DIAGRAM) Consortium et al., 2014]. Across established loci for three of these diseases (no genome-wide significant loci were reported for major depressive disorder), the direction and relative size of effects were consistent between our adjusted ORs and those reported previously (0.92 < Pearson’s *r* < 0.97), though the adjusted ORs were slightly underestimated, with regression slopes between 0.66 and 0.92 (Figure 3). We observed significant (*P* < 0.01) genetic correlations (*r*_*g*_, the proportion of variance in disease liability that is shared between two phenotypes) between our GWAX results and the published GWAS summary statistics for coronary artery disease (*r*_*g*_ = 0.93), major depressive disorder (*r*_*g*_ = 0.67), type 2 diabetes (*r*_*g*_ = 0.91) and Alzheimer’s disease (*r*_*g*_ = 0.44).

### Meta analysis

Motivated by these consistent odds ratios, and in an effort to identify additional risk loci, we performed fixed-effects meta-analysis combining our proxy-case-control association summary statistics with those from the previously published GWAS. This approach implicated 17 novel risk loci at genome-wide significance associated with Alzheimer’s disease, coronary artery disease and type 2 diabetes (Table 1, Figure 4, Figure S4-S6).

Among the novel loci for Alzheimer’s disease include genes invovled in immune surveillance *(SPPL2A,* signal peptide peptidase like 2A) and major histocompatability complex class II signal transduction *(SCIMP,* SLP adaptor and SCK interacting membrane protein) [Friedmann et al., 2004], further highlighting the role of the innate immune system in Alzheimer’ diseasexs etiology [Chan et al., 2015; Gjoneska et al., 2015]. For coronary artery disease, one novel locus resides in an intron of *FGD5* (FYVE, RohGEF and PH domain containing 5), a member of the FGD family of guanine nucleotide exchange factors. *FGD5* has been shown to regulate *VEGFA* (vascular endothelial growth factor) [Kurogane et al., 2012], a key cytokine in the formation of new vessels and potential therapeutic target for heart disease [Taimeh et al., 2013]. For type 2 diabetes, we identified a novel locus in *P1TPNC1* (phosphatidylinositol transfer protein, cytoplasmic 1), a member of the phosphatidylinositol transfer protein family and has been show to be involved in lipid transport between membrane compartments [Garner et al., 2012].

**Table 1.**
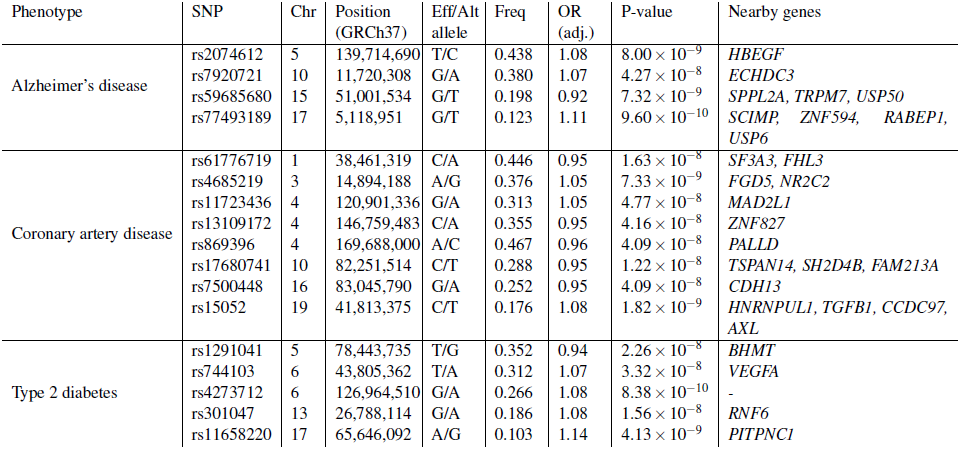
Novel genome-wide significant risk loci identified through proxy-case-control analysis and meta-analysis with published genome-wide association studies of Alzheimer’s disease, coronary artery disease and type 2 diabetes.

**Figure 3.**
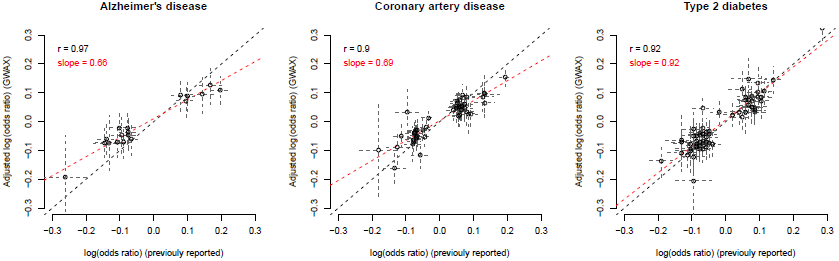
Comparison of adjusted ORs and previously reported case-control ORs at established risk loci for three diseases with publicly available summary statistics. Each point represents a previously reported risk variant and its corresponding effect size. The dashed gray lines are 95% confidence intervals. The dashed red line (and corresponding slope) is the fitted line from least squares regression. The dashed black line is is y = x. Reported effect sizes and list of established risk loci were obtained from - Alzheimer’s disease: Lambert et al. [2013]; coronary artery disease: The CARDIoGRAMplusC4D Consortium [2015]; type 2 diabetes: DIAbetes Genetics Replication And Meta-analysis (DIAGRAM) Consortium et al. [2014].

To further illustrate the utility of the GWAX approach, we also performed case-control GWAS in the UK Biobank (taking case status from medical records) for coronary artery disease (5,685 cases, 109,347 controls) and type 2 diabetes (2,463 cases, 112,273 controls), the results of which were combined with the previously published summary GWAS statistics described above. Of the eight novel coronary artery disease risk loci identified in the GWAX meta-analysis, only two were genome-wide significant in the GWAS metaanalysis. Similarly, none of the five novel type 2 diabetes loci exceeded genome-wide significance in the GWAS meta-analysis (Table S4). These results further demonstrate that using proxy-cases is more powerful than cases for identifying risk loci in population cohorts.

## Discussion

This study demonstrates proof of principle that complex disease risk loci can be identified using the genotypes of unaffected individuals and the phenotypes of their affected relatives. We applied the GWAX approach to 12 common diseases in 116,196 individuals from the UK Biobank and combined our results with publicly available GWAS summary statistics for four of these diseases. We replicated known risk loci and identified 17 novel risk loci at genome-wide significance associated with Alzheimer’s disease, coronary artery disease and type 2 diabetes.

**Figure 4.**
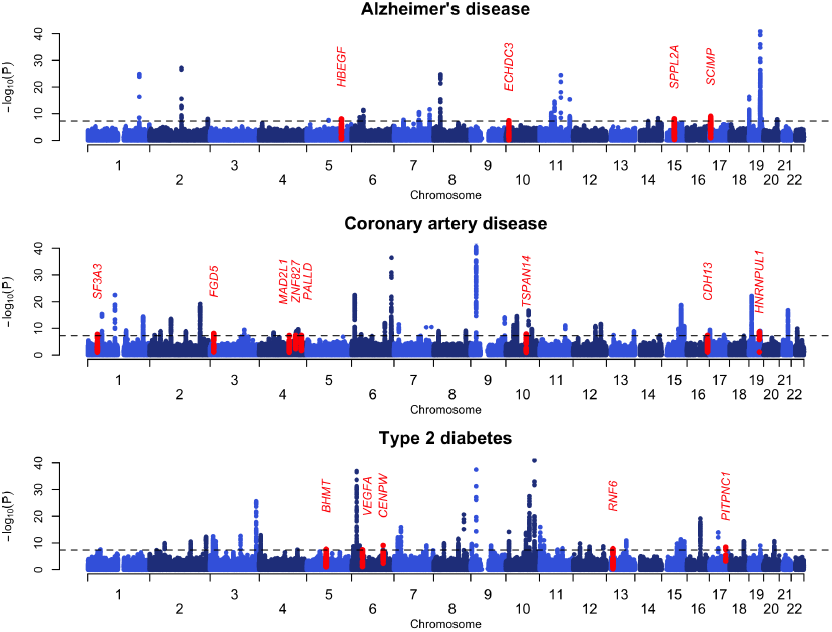
Manhattan plots of fixed-effects meta-analysis results for Alzheimer’s disease, coronary artery disease and type 2 diabetes. Chromosome and positions are plotted on the x-axis. Strength of association is plotted on the y-axis. Novel risk loci are indicated in red. The dashed horizontal line indicates the genome-wide significant threshold of *P* < 5 × ^108^. — ^log^10 P-values are truncated at 40 for illustrative purposes.

Large population cohorts such as the UK Biobank and NIH Precision Medicine Initiative along with participant-driven projects [Dolgin, 2010; Eriksson et al., 2010] are emerging as valuable resources in biomedical research. By performing association mapping using the family members of affected individuals, we partly overcome the ascertainment limitations inherent in these studies.

Future expansions of these approaches may take into account more distant relatives in a formal way, allowing for the phenotypes of all known relatives to be accounted for and analyzed in conjunction with directly genotyped individuals. Genetic studies of complex disorders may progress beyond simple “case” and “control” phenotypes, and instead leverage multiple layers of information into a direct estimate of disease liability. Large crowd-sourced family trees [Ledford, 2013] along with reported phenotypes, demographics, lifestyle surveys, medical records and epidemiological information can be combined to provide robust estimates of both the genetic and environmental components of disease liability [Campbell et al., 2010]. Using liability as a phenotype can also account for ascertainment biases of case-control studies [Hayeck et al., 2015; Weissbrod et al., 2015], and allow for much greater power to identify disease susceptibility variants.

## Methods

### Power calculations

We performed power calculations comparing a study design using true cases and controls to one with proxy-cases and controls, and estimated the sample sizes of each such that power to detect association is equivalent. We also considered the situation where both cases and proxy-cases are available in the context of a population cohort study, where the expected number of cases and proxy-cases depends on disease prevalence and heritability on the liability scale. Details of the power calculations are described in the Supplementary Note.

### UK Biobank data collection

The UK (United Kingdom) Biobank is a large population-based study of over 500,000 subjects aged 40-69 years recruited from 2006-2010 [Sudlow et al., 2015]. Participants entered information about their family history of disease by answering three questions: 1) “Has/did your father ever suffer from?”, 2) “Has/did your mother ever suffer from?”, and 3) “Have any of your brothers or sisters suffered from any of the following diseases?”. Participants were asked to choose among 12 conditions (heart disease, stroke, high blood pressure, chronic bronchitis/emphysema, Alzheimer’s disease/dementia, diabetes, Parkinson’s disease, severe depression, lung cancer, bowel cancer, prostate cancer and breast cancer) and were allowed to select more than one condition. Participants were also given the choice of entering “Do not know”, “Prefer not to answer” or “None of the above”. Throughout this manuscript, we denote heart disease, severe depression and diabetes to refer specifically to coronary artery disease, major depressive disorder and type 2 diabetes, respectively. Case/control statuses for the participants themselves were available via health records (ICD10 diagnoses, Table S5). The UK Biobank received ethics approval from the National Health Service National Research Ethics Service (Ref 11/NW/0382).

### Effective sample size comparisons

For each of the 12 phenotypes, we converted the observed number of cases (or proxy-cases) and controls into an effective sample size (*N_eff_*). The effective sample size is the total sample size where there is an equal number of cases (or proxy-cases) and controls that gives the equivalent power to detect association as the observed unequal sample size. The test statistic for a standard 2 × 2 1-df chi-square test when the number of cases and controls differ is:

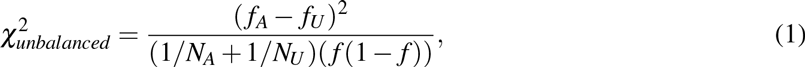

where *N*_*A*_ is the number of cases (or proxy-cases), *N*_*U*_ the number of controls, *f*_*A*_ is the allele frequency in cases (or proxy-cases), *f*_*U*_ the allele frequency in controls and *f* is the overall allele frequency. Under a balanced design where 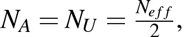, the test statistic becomes:

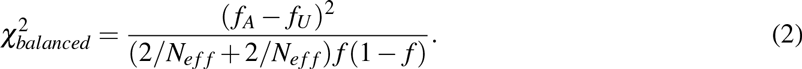

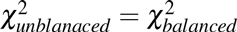 and solving for *N*_*eff*_, we have the effective sample size as a function of observed number of cases (or proxy-cases) and controls:

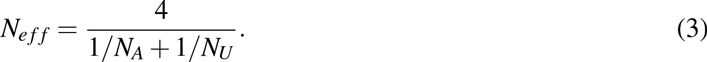

When we report the effective sample size in proxy-cases and controls, we divide *N*_*eff*_ by four to account for power, enabling a direct comparison with the effective sample size when using cases and controls (Supplementary Note).

### Genotyping, imputation and quality control

The UK Biobank May 2015 Interim Data Release included directly genotyped and imputed data for 152,529 individuals. Around 90%, of individuals were genotyped on the Affymetrix UK Biobank Axiom array, while the remainder were genotyped on the Affymetrix UK BiLEVE array. The two platforms are similar with > 95% common marker content (847,441 markers in total). Markers were selected on the basis of known associations with phenotypes, coding variants across a range of minor allele frequencies, and content to provide good genome-wide imputation coverage in European populations for variants with minor allele frequencies > 1%. Genotyped individuals were phased using SHAPEIT2 [Delaneau et al., 2014] and then imputed with the IMPUTE2 [Howie et al., 2009] algorithm using a reference panel consisting of 12,570 haplotypes from a combined UK10K [The UK10K Consortium, 2015] and 1000 Genomes Phase 3 dataset [The 1000 Genomes Project Consortium, 2015]. In total, 73,355,667 polymorphic variants were successfully imputed. Additional information on the genotyping array, sample preparation and quality control can be found in the documents the URLs section. After QC, We took forward 116,196 unrelated individuals of European descent for analysis.

### Genome-wide association by proxy

The genome-wide association by proxy study design is an extreme version of approaches that try to impute unknown genotypes in phenotyped individuals based on the genotypes of close relatives, though these approaches either require accurate pedigree information [Gudbjartsson et al., 2008] and/or sparse genotypes (e.g. microsatellites) on which to impute [Burdick et al., 2006]. Our approach is also similar to that of MQLS [Thornton and McPeek, 2007], a method for association testing in related individuals that allows for combinations of known and unknown phenotypes and genotypes. Indeed, when the genotyped individuals are all of unknown phenotype but with the phenotype of one parent available, MQLS and our approach (using Pearson’s *χ*^2^ test) are mathematically equivalent (Johanna Jakobsdottir, personal communication). However, the current implementation of MQLS does not allow for situations where all genotyped individuals have unknown phenotypes, does not scale to large cohorts and genome-wide data, and cannot handle covariates like principal components to account for population structure. By contrast, using standard logistic regression scales easily to large datasets and can handle covariates without issues.

To perform GWAX in the UK Biobank, for each of the 12 common diseases, subjects were considered proxy-cases if they have at least one affected mother, father or sibling. Subjects who answered “Do not know”, or “Prefer not to answer” were removed from the analysis. All other subjects were considered controls. The total number of proxy-cases and controls for each phenotype are listed in Table S1.

Association between genotype and phenotype was performed on best-guess imputed genotypes (allelic likelihood > 0.9, missingness < 10%, minor allele frequency > 0.005) using logistic regression in PLINK2 [Chang et al., 2015]. For all analyses, we included the subjects’ reported sex, age at recruitment and the first four principal components (estimated directly from the post-QC set of UK Biobank individuals) as covariates. We observed modest genomic inflation across the 12 diseases (1.05 < λ < 1.07; Figure S4).

To test whether these inflation is due to population stratification or reflect a true polygenic signal, we performed LD score regression on the summary association statistics using a set of 1.2 million common SNPs from HapMap3 [Bulik-Sullivan et al., 2015b]. The LD Score regression intercepts were between 0.99 and 1.02 (Figure S7), suggesting that the inflation is due to a true polygenic signal. For the 24 lead SNPs identified with *P* < 5 × 10^−8^, we also performed association testing using a linear mixed model implemented in BOLT-LMM [Loh et al., 2015], where genetic relatedness between the UK Biobank was estimated using 623,852 directly genotyped SNPs. P-values were very similar with those from the logistic regression using 4PCs (Table S3). Together, these results suggest that the effects of population stratification were minimal. As such, we did not adjust association statistics using genomic control.

To enable direct comparison of our effect sizes to those from traditional case-control designs (as well as enabling fixed-effect meta-analysis), we calculate odds ratios using the following approximation. For each SNP, let *f_A_* and *f*_*U*_ be the allele frequencies in true-cases and controls respectively, and

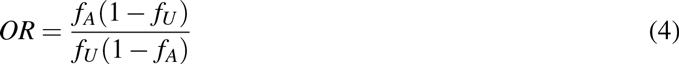

be the true case-control odds ratio. If *f_P_* is the allele frequency in proxy-cases (the vast majority of whom have only one first-degree relative affected with disease), then

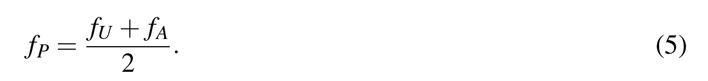

In order to estimate the adjusted odds ratio as a function of the observed allele frequencies in proxy-cases and controls, we substitute *f_A_* into (1):

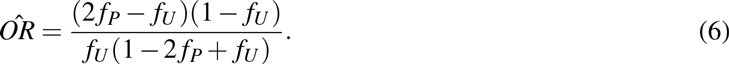

For the range of ORs (< 1.4) typically reported in a GWAS, the log of the adjusted odds ratio derived here is approximately double that of the log odds ratio directly estimated from logistic regression using proxy-cases and controls (Figure S1). As the odds ratios and standard errors from logistic regression take into account covariates, we report adjusted log odds ratios using this doubling approximation rather than directly estimating them from allele frequencies using equation (6). The corresponding adjusted standard error is also double the standard error of the log odds ratio from logistic regression, since 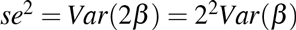.

### Polygenic risk scores

Publicly available GWAS summary association statistics were obtained for Alzheimer’s disease (17,008 cases and 37,154 controls for stage 1 SNPs; plus 8,572 cases and 11,312 controls for 11,632 stage 2 SNPs) [Lambert et al., 2013], coronary artery disease (60,801 cases and 123,504 controls) [The CARDIoGRAM-plusC4D Consortium, 2015], major depressive disorder (9,249 cases and 9,519 controls) [Ripke et al., 2013] and type 2 diabetes (26,488 cases and 83,964 controls) [DIAbetes Genetics Replication And Meta-analysis (DIAGRAM) Consortium et al., 2014].

From these summary statistics, we extracted the reported effect sizes at established loci for Alzheimer’s disease (20 SNPs), coronary artery disease (55 SNPs) and type 2 diabetes (71 SNPs) and constructed polygenic risk scores for each individual in the UK Biobank. No genome-wide significant risk loci were reported for major depression. For a disease with *m* associated SNPs, the polygenic risk score for individual *i* is:

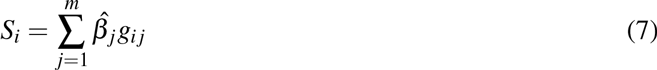

where 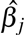 is the reported effect size (log(OR)) of the reference allele of SNP *j* from the previous GWAS, and *g_ij_* is the allele count of the reference allele for individual *i* at SNP *j*. Scores were normalized to mean = 0 and variance = 1. The means of the normalized polygenic risk scores were calculated for groups of individuals who were 1) affected with the disease, 2) unaffected with two affected parents and 3) unaffected with one affected parent. For this last group, the mean risk score was doubled so that it is (in theory) equivalent to the risk score for unaffected individuals with two affected parents. We tested for a significant difference in the mean risk scores for each pair of groups using Welch’s t-test.

### Genetic correlation

For each of the four phenotypes, we estimated the genetic correlation between our GWAX summary statistics and the published GWAS summary statistics using LD score regression with a set of ~ 1.2 million common SNPs from HapMap3 [Bulik-Sullivan et al., 2015a].

### Meta-analysis

Fixed-effects meta-analysis was performed for Alzheimer’s disease, coronary artery disease, major depressive disorder and type 2 diabetes using inverse variance-weighted method for all SNPs that overlap between the publicly available summary statistics and our adjusted odds ratio GWAX results. That is, for each SNP with estimated log odds ratios and standard errors, 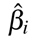and 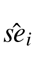 respectively, where *i* = 1 or 2 correponding to the GWAX (adjusted log odds ratio) or GWAS results, the combined effect size is: 
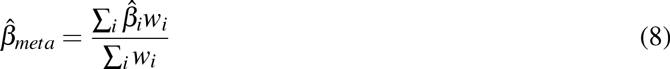

with corresponding standard error and P-value: 
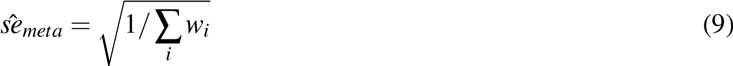

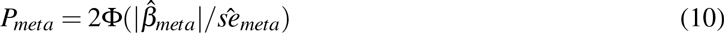

where 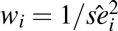 and Φ is the cumulative standard normal distribution.

### Identification of independent risk loci

A locus was considered genome-wide significant if it includes a SNP with association *P* < 5 × 10^−8^. For both the primary proxy-case-control analysis in UK Biobank individuals and meta analyses, independent risk loci were identified using the approximate conditional and joint association method implemented in GCTA (GCTA-COJO). The method performs approximate step-wise conditional association testing using summary association statistics and LD structure from a set of reference genotypes. As such, the SNPs selected from this procedure can be thought to represent the strongest independent signals associated with the phenotype. We ran GCTA-COJO with settings *r*^2^ > 0.9 and *P* < 5 × 10^−8^, and a reference panel consisting of 2,500 randomly selected individuals from the UK Biobank cohort. [Yang et al., 2012].

## URLs

UK Biobank - http://www.ukbiobank.ac.uk

Genotyping and quality control of UK Biobank, a large-scale, extensively phenotyped prospective resource –http://www.ukbiobank.ac.uk/wp-content/uploads/2014/04/UKBiobank_genotyping_QC_documentation-web.pdf

Genotype imputation and genetic association studies of UK Biobank-http://www.ukbiobank.ac.uk/wp-content/uploads/2014/04/imputation_documentation_May2015.pdf

Parkinson’s disease GWAS summary statistics from Nalls et al. [2014] – http://pdgene.org/view?study=1

## Acknowledgement

JZL and JKP are partially supported by the National Institute of Mental Health (NIH grant R01MH106842).We thank Tristan Hayeck and Johanna Jakobsdottir for comments on a draft of this manuscript. This research has been conducted using the UK Biobank Resource. We would like to thank all UK Biobank participants and those involved in sample collection, genotyping, quality control and imputation.

